# The effect of bolus properties on muscle activation patterns and TMJ loading during unilateral chewing

**DOI:** 10.1101/2023.03.02.526627

**Authors:** Benedikt Sagl, Martina Schmid-Schwap, Eva Piehslinger, Hai Yao, Xiaohui Rausch-Fan, Ian Stavness

**Affiliations:** Center of Clinical Research, University Clinic of Dentistry, Medical University of Vienna, 1090 Vienna, Austria; Division of Prosthodontics, University Clinic of Dentistry, Medical University of Vienna, 1090 Vienna, Austria; Department of Bioengineering, Clemson University, 29634 Clemson, SC, United States of America; Department of Oral Health Sciences, Medical University of South Carolina, 29425 Charleston, SC, United States; Department of Computer Science, University of Saskatchewan, SK S7N 5C9 Saskatoon, Saskatchewan, Canada

## Abstract

Mastication is a vital human function and uses an intricate coordination of muscle activation to break down food. Collection of detailed muscle activation patterns is complex and commonly only masseter and anterior temporalis muscle activation are recorded. Chewing is the orofacial task with the highest muscle forces, potentially leading to high temporomandibular joint (TMJ) loading. Increased TMJ loading is often associated with the onset and progression of temporomandibular disorders (TMD). Hence, studying TMJ mechanical stress during mastication is a central task. Current TMD self-management guidelines suggest eating small and soft pieces of food, but patient safety concerns inhibit *in vivo* investigations of TMJ biomechanics.

For this purpose, we have developed a state-of-the-art *in silico* model, combining rigid body bones, finite element TMJ discs and line actuator muscles. To solve the problems regarding muscle activation measurement, we used a forward dynamics tracking approach, optimizing muscle activations driven by mandibular motion. We include a total of 256 different combinations of food bolus size, stiffness and position in our study and report kinematics, muscle activation patterns and TMJ disc von Mises stress.

Computed mandibular kinematics agree well with previous measurements. The computed muscle activation pattern stayed stable over all simulations, with changes to the magnitude relative to stiffness and size of the bolus. Our results agree with the clinical guidelines regarding bolus modifications as smaller and softer food boluses lead to less TMJ loading. The results help to strengthen the confidence in TMD self-management recommendations, potentially reducing pain levels of patients.

## Introduction

The human mandible is able to move intricately using two 6 degree of freedom joints, namely the temporomandibular joints (TMJ). Mastication is a primary function of the jaw region and is vital for human survival^1^. A masticatory cycle uses a complex coordination of activation of jaw opening and closing muscles to break down food. Chewing is generally seen as the orofacial functional task with the highest muscle forces, which in turn might lead to increased temporomandibular joint loading^2^. Since increased TMJ loading is commonly connected with the onset and progression of temporomandibular disorders (TMD), the investigation of TMJ mechanical stress during chewing is an highly relevant task^3,4^. Detailed *in vivo* investigations of TMJ biomechanics are virtually impossible due to the patient health risks caused by the small size and intricate build of the joint. Therefore, *in silico* studies remain the most promising tool for gathering a better understanding of TMJ biomechanics. The objective of the presented study was to investigate the effect of food bolus variables on muscle activation pattern and TMJ disc stress during a unilateral chewing cycle.

Previous *in silico* modeling literature of mastication has mainly focused on multibody modeling, which represents the TMJ as a simplified connection of planar or spline function constraints without a TMJ disc^5–7^. Due to their fast computation times, these simulations are well suited to investigate complex chewing dynamics, but their simplified TMJ representations do not allow for the computation of detailed TMJ loading. Detailed TMJ mechanics are commonly investigated using the finite element (FE) method^8–12^. However, most FE studies of TMJ biomechanics focus on static tasks, like tooth clenching^13–16^ or completely neglect the food bolus^12^. A previous study investigated the effect of bolus size, position and stiffness on TMJ mechanical loading during a dynamic unilateral chewing cycle using a combined multibody-FE model and a linear elastic food bolus^17^. While this study gave valuable insight into the effect of the bolus on TMJ loading it used the same muscle activation pattern for all simulations. Parameters like bolus stiffness and size are expected to have an influence on muscle activation, which consequently could have an additional effect on TMJ loading^2^. Moreover, *in vivo* measurement of masticatory muscle activation is hard, due to the small size and overlapping of chewing muscles and masticatory EMG is generally only performed for the masseter and anterior temporalis^18^. We previously tried solving this problem by means of a state of the art forward dynamics tracking approach to optimize muscle activation from a mandibular motion target input^19^.

A possible influence of food bolus properties, such as stiffness and size of the food piece, on TMJ biomechanics is of special interest, since clinicians often recommend a diet consisting of small and soft pieces of foods as part of TMD self management guidelines^20–23^. While these recommendations sound reasonable, only a rather limited amount of scientific evidence exists^24,25^. Consequently, a proper understanding of the effect of food bolus properties on muscle activation patterns and in turn TMJ loading could help inform and update TMJ self management guidelines. In fact, recently Jurt et al. 2020 reported a potential effect of large bolus sizes on TMJ loading via dynamic stereometry^26^.

Previous investigations of chewing cycle biomechanics were mostly performed using “dynamic stereometry”, which investigates the distances between the mandibular condyle and articular eminence^26–30^. Increased mechanical TMJ loading suggests a decrease in joint space due to compression of the TMJ disc. Using this method the group has found a smaller joint space on the non-chewing side TMJ as well as during the closing portion of the chewing cycle. These findings would indicate that chewing on the painful side would be beneficial to minimize mechanical loading on the painful TMJ. While dynamic stereometry can give important *in vivo* insight in mandibular kinematics during chewing, mechanical loading can only be inferred.

Overall, our study aims to investigate the effect of bolus stiffness, bolus position as well as bolus size on muscle recruitment and consequently TMJ loading during a dynamic, unilateral chewing cycle. We use innovative *in silico* approaches, like our combined rigid body – finite element model, driven by line actuators, as well as a forward dynamics tracking approach to simulate muscle driven, dynamic chewing tasks. For this purpose, a total of 256 simulations were performed and computed mandibular motion, muscle activation as well as TMJ disc stress were analyzed. The knowledge gained during the course of this study has potential impact on future self-management guidelines for TMD.

## Results

### Mandibular Kinematics

Of the 256 simulations 231 finished successfully. As expected, simulations with very large and stiff boluses tended to overload the FE discs, which lead to the simulations failing.

Generally speaking, mandibular motion successfully followed the expected teardrop shaped ICP motion pattern (Figure 1). Bolus stiffness and size had some impact on mandibular kinematic due to differences in computed muscle activation patterns. Jaw opening was successfully scaled to open the mouth to fit the bolus in between the jaws without any excessive gape. The maximum displacement of the midincisor point was 3.82±1.27mm in the mediolateral direction, 6.08±3.0278mm in the anteriorposterior direction and -13.07±6.41mm in the inferiorsuperior direction. The maximal displacement of the chewing side condyle was 0.63±0.57mm in the mediolateral direction, -0.53±0.69mm in the anteriorposterior direction and -0.82±0.89mm in the inferiorsuperior direction. For the non-chewing side condyle, the maximum displacements were - 1.01±0.57mm in the mediolateral direction, -2.7±0.49mm in the anteriorposterior direction and - 2.72±0.62mm in the inferiorsuperior direction.

**Figure 1:**
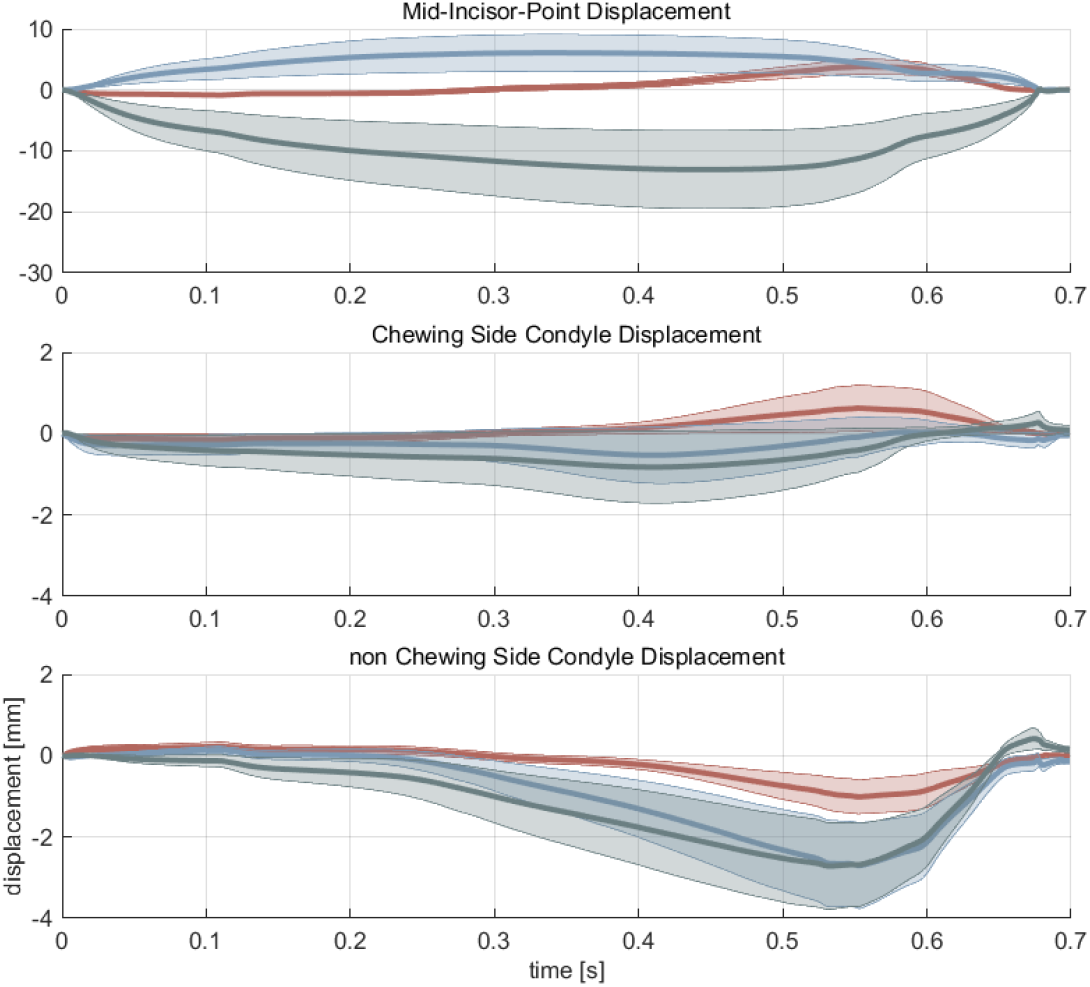
Displacement [mm] vs. time [s] for the mid-incisor point (top), chewing side condyle (middle) and non-chewing side condyle (bottom). Red: mediolateral direction (positive values mean towards the chewing side); Blue: anteriorposterior direction (negative values mean anterior); Green: inferiorsuperior direction (negative values mean inferior). The solid line represents the mean displacement across all simulations. The filled area denotes ± one standard deviation.

### Muscle activations

Comprehensive heat maps of the mean activation of all included muscles during the opening and closing part of the chewing cycle for all included simulation set-ups can be found in the supplemental material (Supplemental Figure 1-4). Overall, the muscle activation pattern stayed relatively stable, with respect to time, throughout all simulations, with only minor differences in pattern. Figure 2 presents horizon graphs for four selected examples, moving bolus position and changing bolus stiffness.

**Figure 2:**
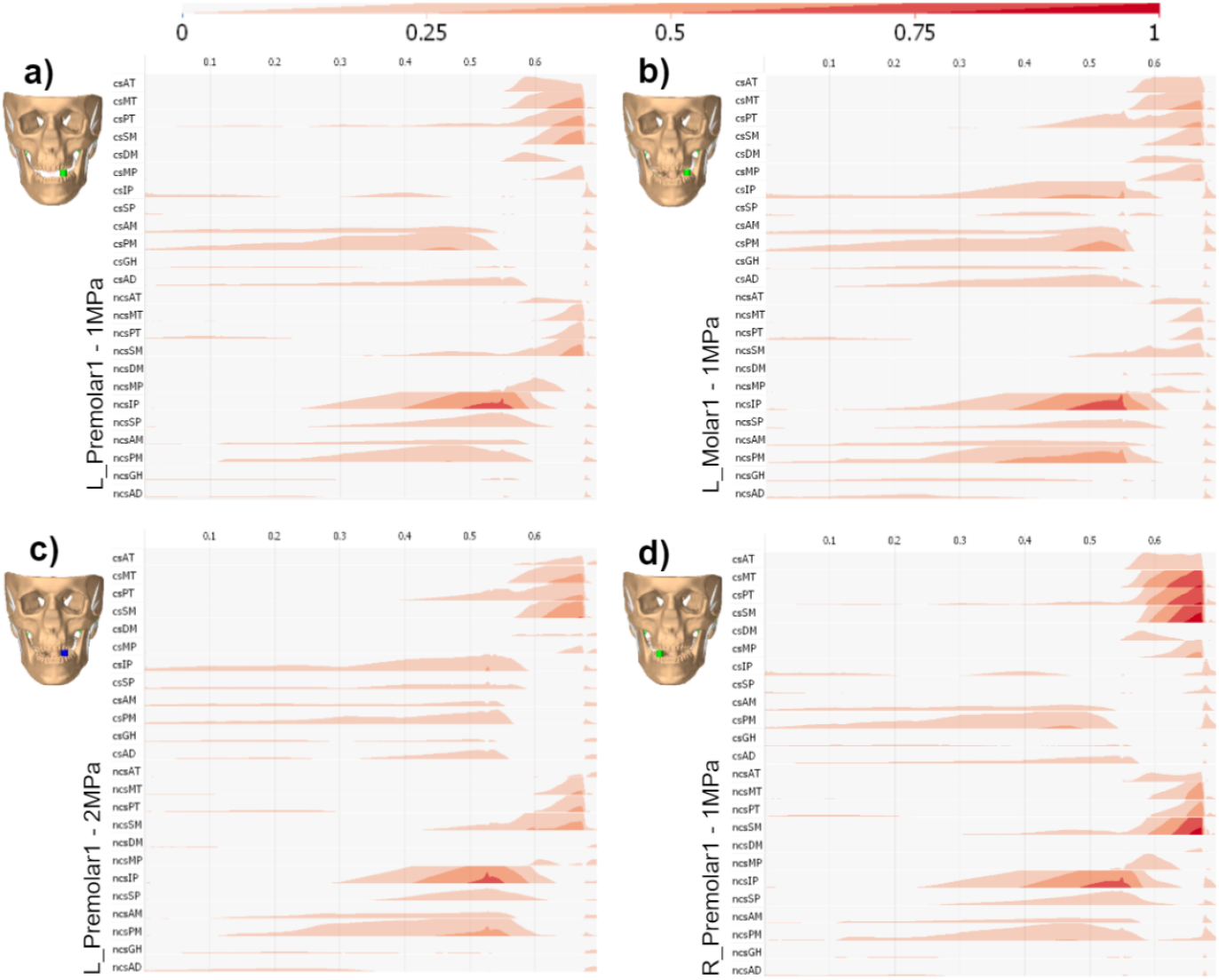
Horizon graphs for representative examples of muscle activation pattern changes. a) chewing on left first premolar with a 10mm bolus with a stiffness of 1MPa; b) schewing on left first molar with a 10mm bolus with a stiffness of 1MPa; c) chewing on left first premolar with a 10mm bolus with a stiffness of 2MPa; d) chewing on right first premolar with a 10mm bolus with a stiffness of 1MPa

Additionally, scaling of muscle forces (more opening muscle activation for larger and/ or more posterior boluses and more closing muscle force for larger and/ or stiffer boluses) could be seen. The opening movement is generally driven by activating the chewing side lateral pterygoid muscle, with higher activations seen in the inferior head, mylohyoid muscle, geniohyoid muscle and anterior digastric muscle. On the non-chewing side, a strong activation of the lateral pterygoid muscle can be seen together with mylohyoid muscle, geniohyoid muscle and anterior digastric muscle activation. Additionally, a small bilateral activation of the posterior part of the temporalis muscle was computed. During closing, mostly activation of the temporalis muscle, masseter muscle and medial pterygoid muscle was observed. Activation of these muscles generally occurs bilaterally, with higher activations reported for the chewing side.

### TMJ disc stress

Figure 3 shows the mean, over all finished simulations, von Mises stress for the chewing and non-chewing side discs over the course of the full chewing cycle. The mean stress was higher on the non-chewing side compared to the chewing side. Additionally, higher TMJ disc stress can be seen during the closing portion of the chewing cycle for both discs. Figure 4 and 5 show heat maps of the mean von Mises stress, over the 200 maximally loaded nodes, for both discs during the closing portion of the chewing cycle.

**Figure 3:**
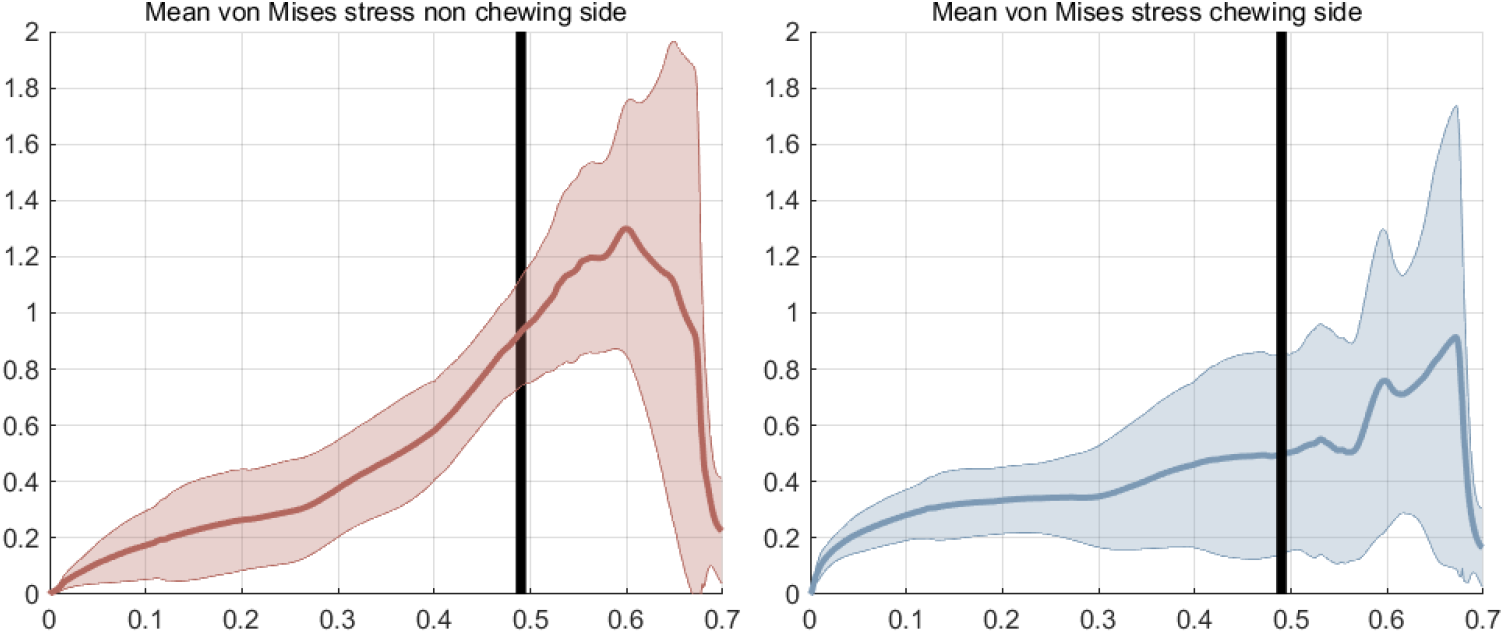
Mean von Mises stress over all included simulations for the non-chewing side (red) and chewing side (blue) TMJ discs; solid, colored line represents the mean stress across all simulations; filled area denotes ± one standard deviation; black vertical line represents border between opening and closing part of chewing cycle.

**Figure 4:**
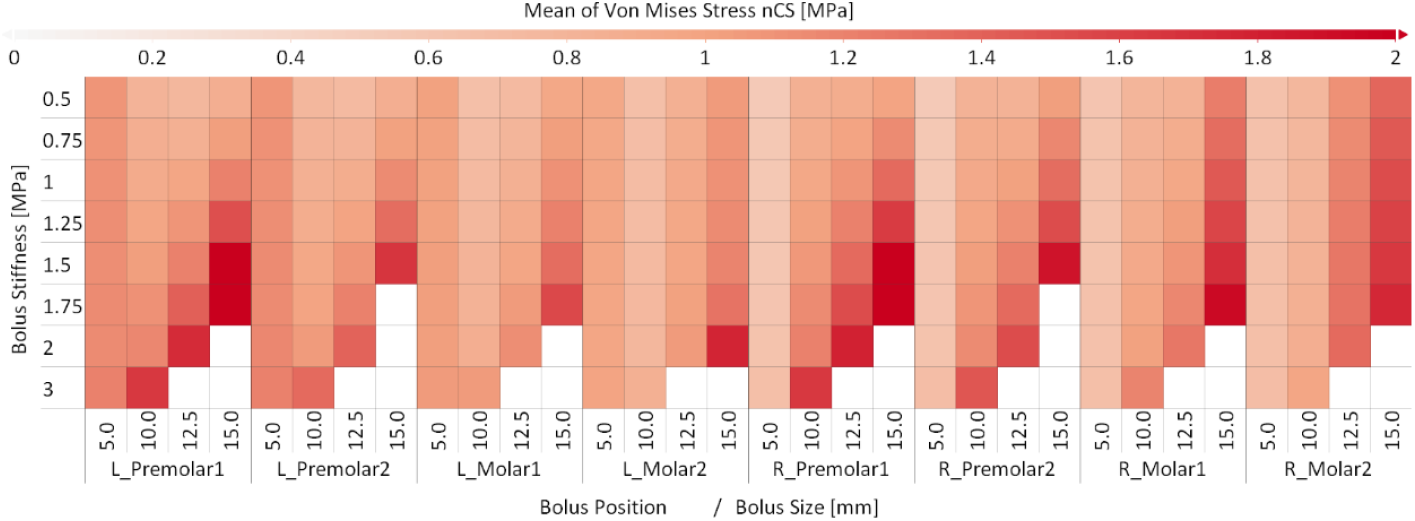
Mean von Mises stress of the 200 highest loaded nodes on the non-chewing side disc for all simulations. L_Premolar1: left first premolar; L_Premolar2: left second premolar; L_Molar1: left first molar; L_Molar2: left second molar; R_Premolar1: right first premolar; R_Premolar2: right second premolar; R_Molar1: right first molar; R_Molar2: right second molar.

**Figure 5:**
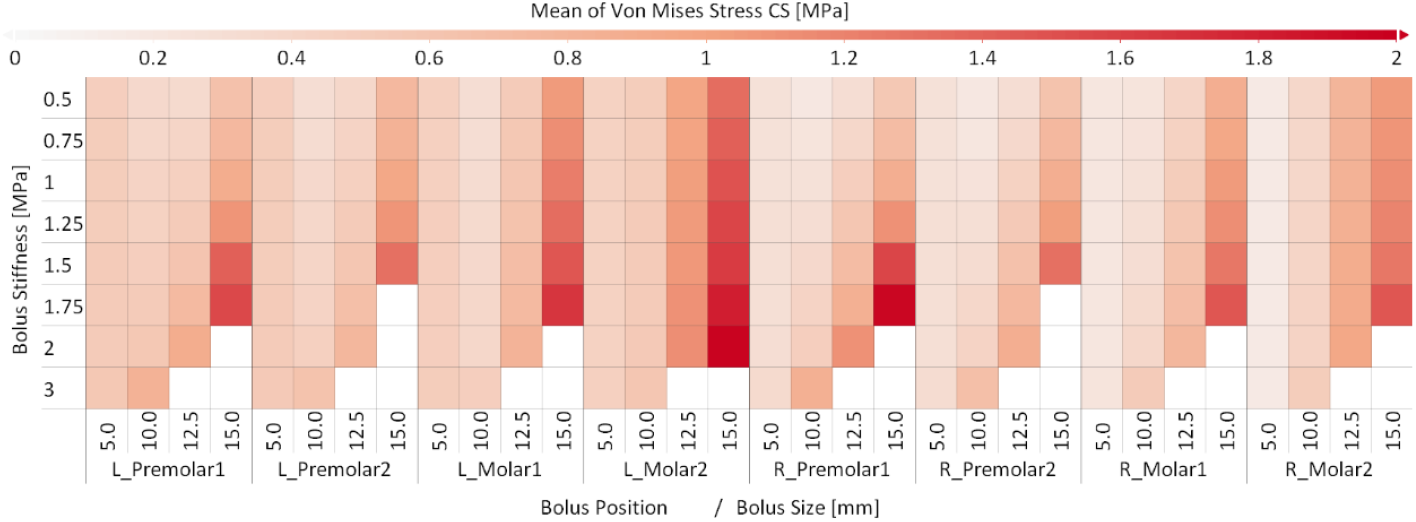
Mean von Mises stress of the 200 highest loaded nodes on the chewing side disc for all simulations. L_Premolar1: left first premolar; L_Premolar2: left second premolar; L_Molar1: left first molar; L_Molar2: left second molar; R_Premolar1: right first premolar; R_Premolar2: right second premolar; R_Molar1: right first molar; R_Molar2: right second molar.

Again, non-chewing side disc stress was generally higher compared to the chewing side disc. This holds true for all combinations except for simulations on the second molar using the largest 15mm bolus. The highest values were seen with 2.56 MPa for chewing on the left first premolar with a 15mm bolus and a stiffness of 1.75 MPa on the non-chewing side disc and 1.97 MPa for chewing with a 2MPa stiff and 15mm large bolus on the left second molar.

## Discussion

Our study offers an interdisciplinary and innovative investigation into the detailed biomechanics of the TMJ during a dynamic, muscle driven unilateral chewing cycle. A total of 256 simulations were performed to thoroughly investigate the potential effect on bolus size, position and stiffness on TMJ mechanical loading. Overall, our mandibular motion and muscle activation results agree well with previous EMG and jaw tracking studies and our findings with regards to mechanical TMJ stress fit well into and augment the current literature. Overall, this paper presents an *in silico* tool for the assessment of chewing biomechanics, a task virtually impossible to achieve *in vivo*. The presented results are of high relevance for the better understand of the complex masticatory region and could help improve and adapt current clinical guidelines for TMD self-management.

The simulations all computed realistic chewing motions, following the characteristic teardrop shape of the interincisal point. Maximum mouth opening, mediolateral and anterior posterior excursion lie within the range reported in literature^5,31^. Since condylar motion has a large influence on TMJ loading a correct representation of this factor is of utmost importance. Our values for maximum excursion of 0.63±0.57mm in mediolateral direction, -0.53±0.69mm in anterioposterior direction and -0.82±0.89mm in inferiorsuperior direction for the chewing side and -1.01±0.57mm in mediolateral direction, -2.7±0.49mm in anteriorposterior direction and -2.72±0.62mm in inferiorsuperior direction for the non-chewing side are relatively close to reported literature values of 0.5±0.6mm, 0.7±0.7mm and 3.1±1.5mm respectively for the chewing and 0.3±0.5mm,3.5±1.6mm and 5.3±2.0mm respectively for the non-chewing side^32^. These comparisons with previous literature show that our simulations were able to compute a realistic chewing motion for all bolus positions, sizes and stiffness values.

Interestingly, all simulations used a comparable muscle activation pattern (Figure 2). The opening cycle was characterized by submental and lateral pterygoid muscle activation with higher activation of the posterior mylohyoid muscle and the inferior head of the lateral pterygoid muscle. The closing phase facilitated activations of all closing muscles with highest activations for the medial and posterior parts of the temporalis and the superficial masseter on the chewing side as well as the posterior part of the non-chewing side temporalis. Activation of the lateral pterygoid muscle, especially on the non-chewing side, can also be seen during the closing phase of the simulation. This is most likely due to the fact that the non-chewing side lateral pterygoid is used to pull the condyle forwards to achieve an appropriate level of lateral mandibular motion, without producing non physiological lateral shifts of the mandible. While the muscle activation pattern stays relatively stable, with regards to muscle activation timing, the magnitude of activation is clearly governed by stiffness and size of the bolus, while position seems to have a diminishing effect, except for large boluses on posterior teeth. When comparing the muscle activation pattern to previous EMG and computational simulations studies^5,33–36^ a good agreement can be seen, with differences mostly in magnitude, but an overall well-fitting activation pattern. This is especially interesting, since no *a priori* knowledge of muscle activation is present in the current model and the muscle activation pattern is solely based on the quadratic optimization performed by the tracking algorithm. This fact further increases confidence in the computed results.

Bolus position seemed to play a lesser role in terms of TMJ loading. Only chewing on the second molar with the largest included bolus of 15mm lead to increased loading. We think that this is caused by the relatively large mouth opening that is necessary to chew through such a thick bolus at this posterior position. Moreover, since the speed of the chewing cycle is kept at 0.7s for all trials, the simulations with very large mouth opening have to open the mouth faster, which in turns means larger muscle activation and in turn potentially an increase in TMJ loading. This trend is especially apparent on the chewing side disc, which is generally loaded less during the chewing cycle. We previously found a decrease of TMJ stress for more posterior chewing positions when keeping the muscle activation stable^17^. This effect diminished in the current study. We think that by allowing the muscle activation pattern to change, more posterior tooth positions generally computed lower magnitudes of closing muscle activation. This modulation of necessary closing force potentially negates the previously seen effect of bolus position on TMJ loading, when keeping the muscle activation pattern the same.

In agreement with previous literature, the stress was higher on the non-chewing side and during the closing portion of the masticatory cycle^26,28,37^. As expected, an increase in bolus stiffness lead to an increase in mechanical loading of the TMJ. This is most likely caused by the increase in activation of the closing muscles to collapse the bolus. This observation is consistent with previous modeling studies^17^ as well as investigations of intra articular space^28^. Moreover, literature reported a reversal of this effect for very large (15mm) boluses, with softer boluses computing higher von Mises stress and a smaller intraarticular space^17,26^. However, we did not see this trend in our simulations. Both above-mentioned studies used boluses that were too stiff to fully bite through them, which might have led to the aforementioned behavior. We previously theorized that a softer bolus, which is still too hard to bite through, allows for more compression and hence brings the mandible closer to the maxilla. This leads to a more optimal position for closing force creation. In our study, all simulations successfully bit through the bolus, which would negate this effect. A clear increase in TMJ loading can be seen when increasing bolus size. This result is consistent with our previous findings^17^ and makes sense since the current simulation models are using a bolus that acts like a “smoothed linear spring”. This leads to the fact that more force is needed to chew through a larger bolus when keeping the material stiffness the same and consequently the higher muscle force will increase TMJ stress.

Overall, our results support the common suggestions to chew on the painful side to reduce mechanical loading. Additionally, our study suggests that the previously reported effect of bolus position on mechanics, when keeping the muscle activation stable, disappears due to the differences in closing muscle force. Bolus size and stiffness both had a clear effect on TMJ loading, which support the self-management guidelines of eating soft and small pieces of food to limit TMJ stress and consequently potentially decrease joint pain. The presented clinical implications have to be applied carefully and clinical trials to verify their results are still necessary.

In previous work the bolus was modeled as a linear elastic spring that increases in force with decreasing distance between the upper and lower teeth until it collapses right before tooth contact^5,17^. While this is an acceptable simplification for forward simulations (with predefined muscle activation patterns), the fact that the bolus force jumps from a high value to zero in a single time step leads to stability issues when combined with our muscle activation optimization method. Preliminary tests showed that the simulations tended to go unstable right at, or shortly after, bolus collapse. At this point in time high muscle forces were computed by the model, since the bolus was applying its highest force, and then the bolus collapsed and applied a force of zero. This change happens in a single time step, which leads to the model trying to recover from this immediate change by using large, short muscle activation spikes, causing a multitude of simulations to fail. Hence, we decided to use a Cubic Hermite spline to improve the bolus force behavior with respect to the mouth gap (Figure 6c). The applied function still has a relatively steep decline in force once the teeth get very near to each other, but the function is differentiable and no spikes in muscle activation have been recorded around bolus collapse. Nevertheless, a very small number of simulations still failed around bolus collapse, which implies that there might be merit in developing a more sophisticated bolus model for a future investigation.

**Figure 6:**
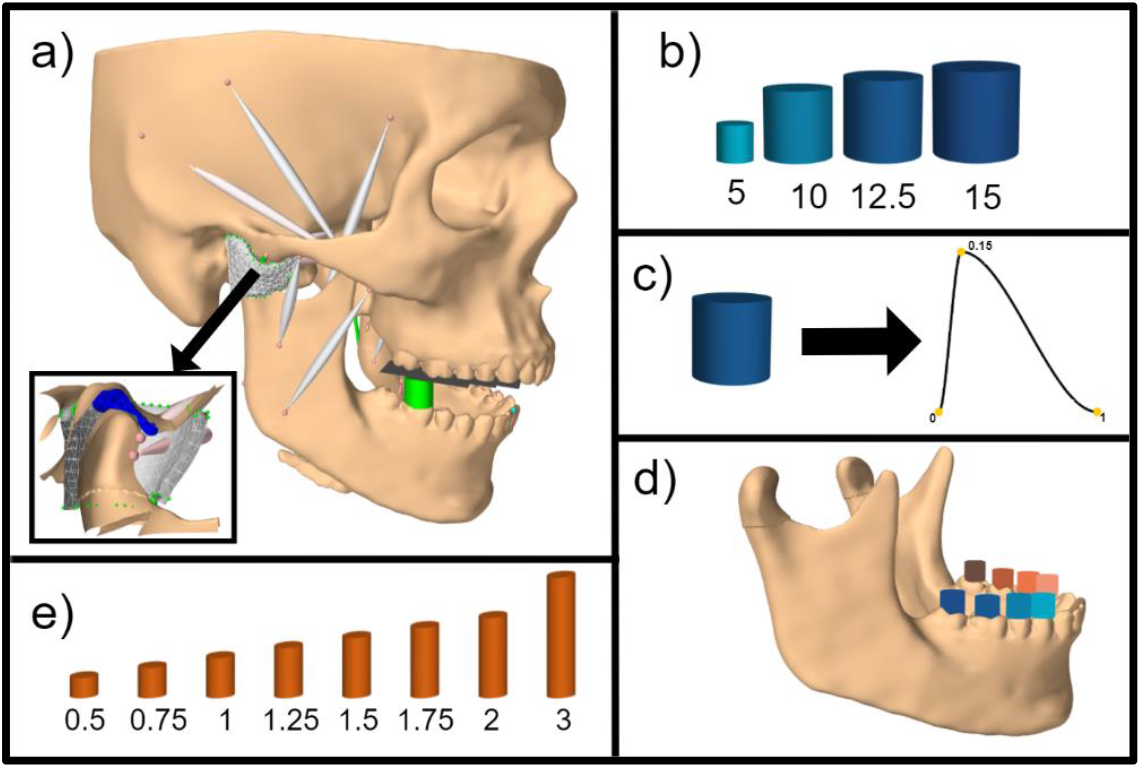
Graphical overview of study; a) model used in study; insert shows cut through TMJ; b: included bolus sizes; c) Hermite spline function used to model bolus force; d) included chewing positions; e) Bolus stiffness values [MPa] included in the study

This *in silico* study includes some modeling assumptions and limitations. First, while this study included a thorough investigation of the effect of bolus variables with a total of 256 tested combinations the model itself was built from the morphology of a single symptom free volunteer. Furthermore, substantial uncertainties with regards to the mechanical properties of soft tissue structures of the orofacial region remain. Additionally, the forward dynamics tracking optimization scheme facilitated to compute the muscle activation patterns involves some assumptions to solve the redundancies related to the muscle force sharing problem^19^. Teeth were simplified using planar external forces, essentially leading to the model using fully flat teeth, which is an oversimplification of the tooth contact behavior. We do think that this problem is negligible for mastication, because the motion is generally performed without tooth contact and the highest joint stress occurs during closing, not during tooth contact. Our investigation includes bolus stiffness values from 0.5 to 3MPa, which roughly spans a range from gouda cheese to raw green turnips^38^. We think this is the appropriate range of values for our representation of the bolus as a “quasi linear” spring facilitating Cubic Hermite splines, since stiffer foods, like nuts, are generally also more brittle and cannot be appropriately modeled using the current bolus setup.

As previously stated, we do think that the presented simulations hold clear merit as comparative investigations of the effect of bolus variables on chewing muscle activation and TMJ loading, but the absolute numerical values should be interpreted with caution. The model currently uses a hyperelastic Mooney Rivlin material model for the TMJ discs, which is commonly used in modeling studies^39–42^, but more complex material models have been suggested to better capture the behavior of the disc^10,43,44^. Besides, *in vivo* measurements of tissue properties for the tissues of the masticatory region are virtually impossible and due to the patient specific modeling approach a rather large amount of anatomical variation is to be expected. Hence, the presented absolute values should not be used as thresholds^45^. They should rather be understood as a comparative analysis of the effect of bolus parameters on muscle activation and TMJ disc loading, which is still of high relevance to better advise clinicians during the decision process with respect to food modifications for TMD patients.

## Material & Methods

The *in silico* model is based on high-resolution data of a symptom free, male volunteer. The institutional review board of the Medical University of Vienna approved the study (Nr. 1190/2017) and written informed consent was obtained. Image acquisition and model development have been previously discussed in detail^46,47^. In short, we used a full skull CT scan together with a full skull MRI scan and high resolution TMJ MRI scans to create surface meshes of all relevant structures and to define muscle paths. The model was created in the ArtiSynth modeling toolkit (www.artisynth.org)^48^. All bones were modeled as rigid bodies and muscles were included as line actuators^49^. The TMJ articular cartilage layers were modeled as elastic foundation contact layers^50^ and the TMJ discs were included as FE bodies, to enable the detailed examination of TMJ disc stress. We used a Mooney Rivlin material with values (C1 = 9 · 10^5^Pa and C2 = 9 · 10^2^Pa) based on previous literature^40^. The TMJ capsule was modeled using approximately 400 FE wedge elements per side and inextensible cables were added to represent the TMJ ligaments^17^. Data visualization was performed using Visplore Discovery (Visplore, Vienna, Austria) and Matlab (The MathWorks, Natick, MA, USA). Figure 6 gives a graphical overview of the study design.

Due to the problems with recording reliable muscle excitation *in vivo*, especially for the deeper chewing muscle, we used a forward dynamics tracking approach to compute the optimal, with respect to the optimization function described below, muscle activation pattern. A detailed theoretical explanation of the forward dynamics tracking algorithm can be found in a previous publication^19^.

In short, the method incorporates mandibular motion in our quadratic optimization function, as follows:

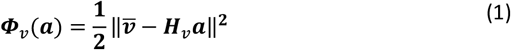

where *Φ*_*ν*_ is the quadratic optimization goal that minimizes the difference between a movement goal, 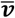, and the mandibular motion created by a set of muscle activations, represented by ***H***_*ν*_*α*. Each row of ***H***_*ν*_ describes the motion of the mandible for the full activation of an individual muscle and is used to speed up calculations.

The optimization equation solved at each time step is:

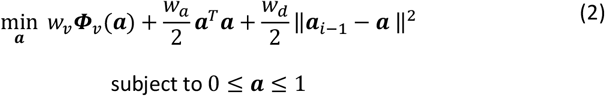

where *w*_*ν*_, *w*_*a*_ and *w*_*d*_ are the respective weights for each cost term (typically *w*_*d*_ < *w*_*a*_ ≪ *w, w*). Since the system is underdetermined, the *l*^2^ norm regularization term 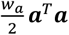 is added. Additionally, we added a damping term ‖*α*_*i*−1_ − *α* ‖^2^ to prevent the solution from oscillating between two possibly feasible states, a behavior that could again be caused by the multitude of muscles in the small area. The weights of the regularization term and damping term were set to 0.0005 and 0.000005, respectively. Motion target for the inter-incisal-point as well as the left and right second mandibular molars were used to drive the chewing motion. Additionally, a static motion target was added to both condyles, which penalized large mediolateral shifts of the mandible^30,32^. Weights for each motion target and directional sub weights were manually optimized. Table 1 presents all sub weights used.

**Table 1:** Sub weights for all included kinematic targets

|  | Mediolateral | Anteriorposterior | Inferiorsuperior |
| --- | --- | --- | --- |
| Interincisal Point | 1 | 0 | 1 |
| Second Molar (Left + Right) | 0 | 0.9 | 0.85 |
| Condyle (Left + Right) | 1 | 0 | 0 |

To create appropriate mandibular input motions, left and right side chewing cycle forward simulations were run using a previously optimized muscle activation pattern^17^. Since the aim of this step was to compute a realistic mandibular motion some additional kinematic constraints were added to keep condylar motion in line with previous recordings^32,51,52^. To accommodate the different sizes of the included boluses mouth opening was scaled, while mediolateral movement was kept stable in accordance with previous literature^53^.

The forward dynamics chewing model previously utilized by our group modeled tooth contact between flat teeth using a combination of unilateral constraints^17^, but unilateral constraints are not compatible with the current quadratic optimization scheme used by our forwards dynamics tracking approach, which only allows bilateral constraints^19^. To circumvent this problem contact between teeth was modeled using an external force component. The force was computed relative to the distance of the maxillary and mandibular tooth rows using a quadratic force gain, a stiffness of 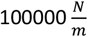 and damping constant of 50. This set up leads to a behavior comparable to a unilateral constraint, since force is only applied when the tooth rows are very close to each other and with a very fast force increase.

The food bolus was previously modeled as a linear spring, which increased the spring force with decreasing distance between the upper and lower jaw^5,17^. This function is non-differentiable at bolus collapse (distance is 0). To improve model stability we replaced the linear function with a one dimensional Cubic Hermite Spline (Figure 6c). For mouth gaps larger than the bolus, size no force was applied and force application starts once the mouth gap reaches the predefined bolus size. The force increases approximately linearly until the mouth gap reaches 1% of the bolus size, at which point the bolus collapses. Table 2 gives an overview of all included bolus variables. All possible combinations of bolus variables were computed to investigate their effect on muscle activation patterns as well as TMJ loading.

**Table 2:** Values of bolus variables investigated in this study

|  | Value |
| --- | --- |
| Bolus Position | first premolar, second premolar, first molar and second molar on both sides |
| Bolus Size | 5mm, 10mm, 15 mm |
| Bolus Stiffness | 0.5 MPa, 0.75 MPa, 1 MPa, 1.25MPa, 1.5MPa, 1.75MPa, 2 MPa, 3 MPa |
| Bolus Position | first premolar, second premolar, first molar and second molar on both sides |

We report mean von Mises stress over the 200 nodes with the highest stress for the chewing and non-chewing side TMJ discs, mean muscle excitation patterns as well as their correlation with TMJ disc loading. The mean stress over the 200 maximally loaded nodes was chosen, because it appropriately represents the region of maximum stress of the disc without being too sensitive to single nodes with extremely high stresses. This approach allows us to investigate the maximal loading on the TMJ disc while keeping our results comparable and stable.

## Acknowledgements

The authors want to thank John Lloyd and the ArtiSynth team for their helpful input regarding *in silico* modeling during the course of the project.

## Conflict of interests

The authors declare no competing interests.

## Contributions

BS, EP, MSS, XRF, HY and IS conceived and designed the study, BS and MSS acquired data, BS, HY, XRF and IS interpreted and analyzed the data, BS drafted the manuscript and all authors revised the manuscript.

## Data Availability

The data presented in this study are available on request from the corresponding author.

---

**Supplemental Figure 1:**
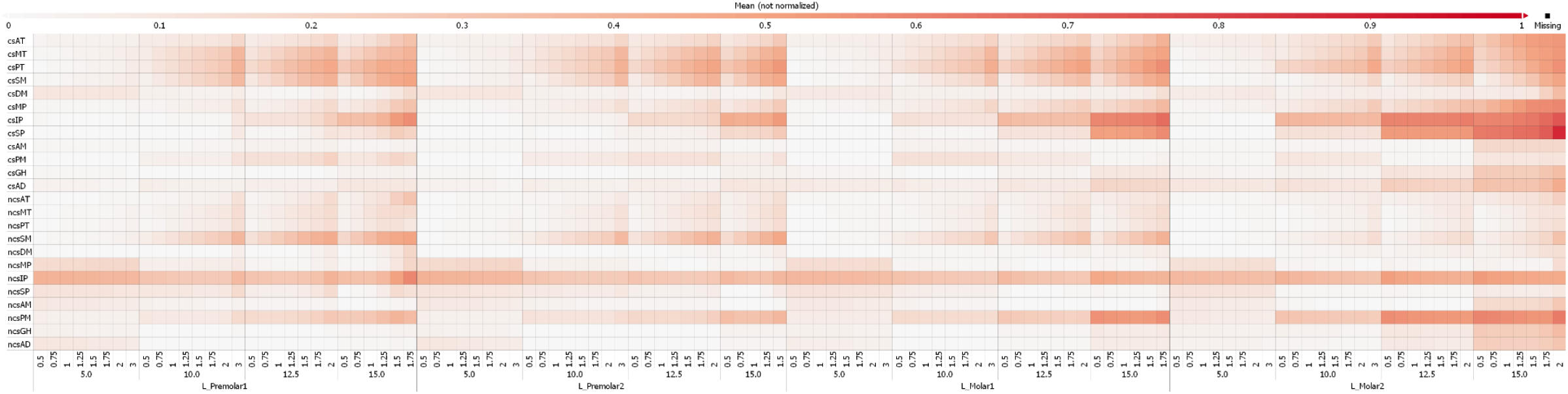
Mean muscle activation for all muscles during closing part of chewing cycle when chewing on the left side; 5, 10, 12.5 and 15 represent bolus sizes in mm, 0.5, 0.75, 1, 1.25, 1.75,2, 3 are bolus stiffnesses in MPa; chewing side (cs) and non chewing (ncs); AT, anterior part of temporalis; MT, medial part of temporalis; PT, posterior part of temporalis; SM, superficial head of the masseter; DM, deep head of the masseter; MP, medial pterygoid; IP, inferior head of lateral pterygoid; SP, superior head of lateral pterygoid; AM, anterior mylohyoid; PM, posterior mylohyoid; GH: geniohyoid; AD: anterior part of the digastric.

**Supplemental Figure 2:**
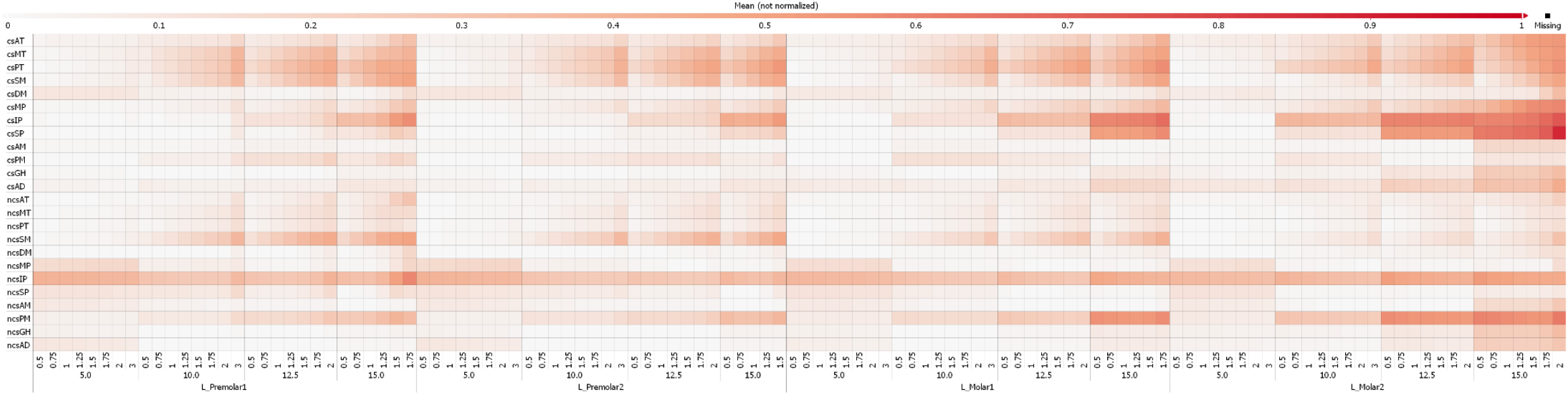
Mean muscle activation for all muscles during closing part of chewing cycle when chewing on the right side; 5, 10, 12.5 and 15 represent bolus sizes in mm, 0.5, 0.75, 1, 1.25, 1.75,2, 3 are bolus stiffnesses in MPa; chewing side (cs) and non chewing (ncs); AT, anterior part of temporalis; MT, medial part of temporalis; PT, posterior part of temporalis; SM, superficial head of the masseter; DM, deep head of the masseter; MP, medial pterygoid; IP, inferior head of lateral pterygoid; SP, superior head of lateral pterygoid; AM, anterior mylohyoid; PM, posterior mylohyoid; GH: geniohyoid; AD: anterior part of the digastric.

**Supplemental Figure 3:**
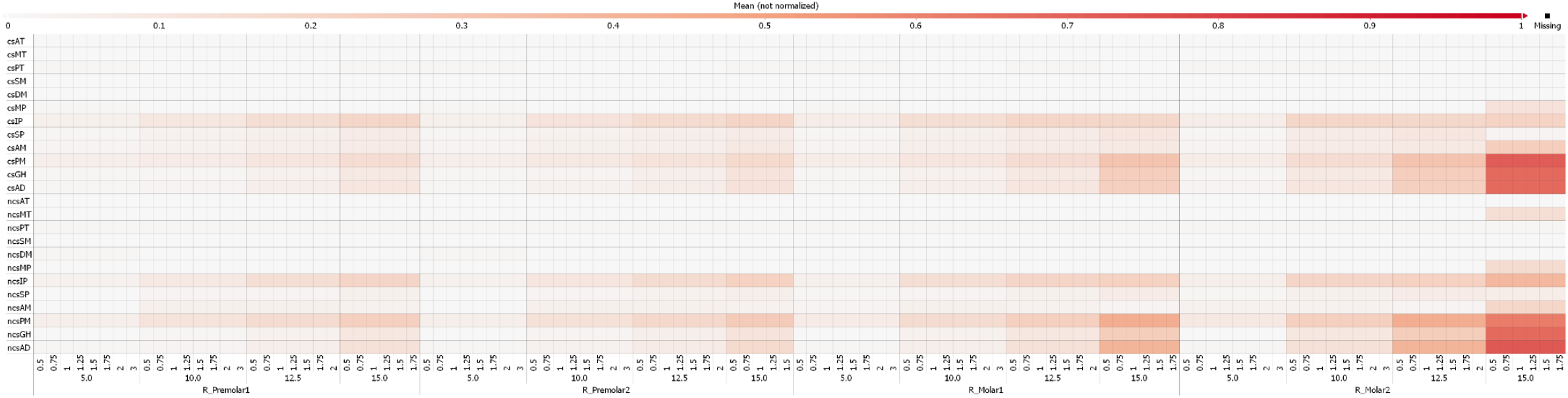
Mean muscle activation for all muscles during opening part of chewing cycle when chewing on the right side; 5, 10, 12.5 and 15 represent bolus sizes in mm, 0.5, 0.75, 1, 1.25, 1.75,2, 3 are bolus stiffnesses in MPa; chewing side (cs) and non chewing (ncs); AT, anterior part of temporalis; MT, medial part of temporalis; PT, posterior part of temporalis; SM, superficial head of the masseter; DM, deep head of the masseter; MP, medial pterygoid; IP, inferior head of lateral pterygoid; SP, superior head of lateral pterygoid; AM, anterior mylohyoid; PM, posterior mylohyoid; GH: geniohyoid; AD: anterior part of the digastric.

**Supplemental Figure 4:**
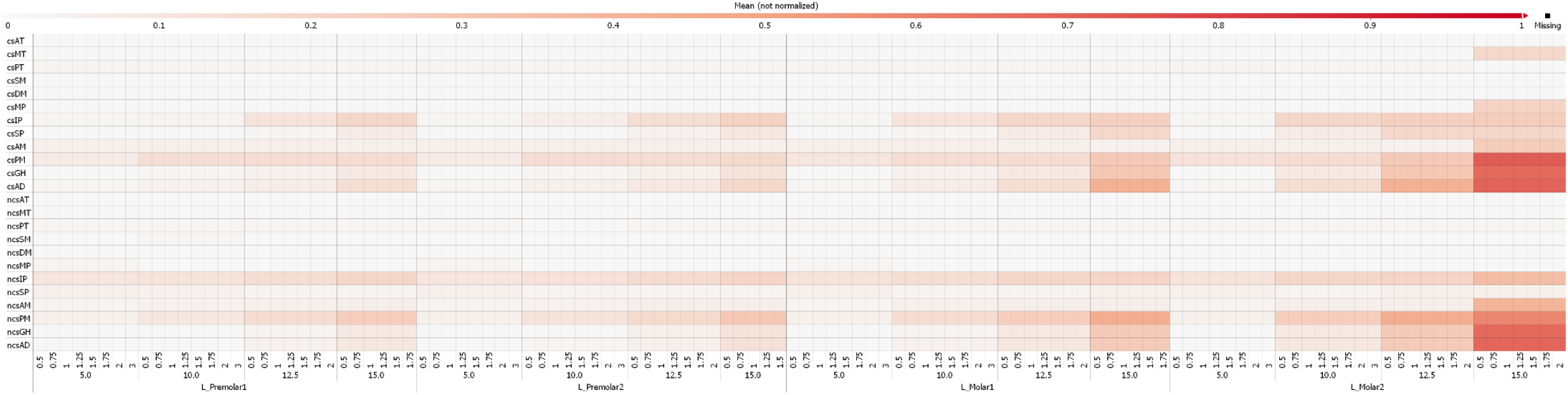
Mean muscle activation for all muscles during opening part of chewing cycle when chewing on the left side; 5, 10, 12.5 and 15 represent bolus sizes in mm, 0.5, 0.75, 1, 1.25, 1.75,2, 3 are bolus stiffnesses in MPa; chewing side (cs) and non chewing (ncs); AT, anterior part of temporalis; MT, medial part of temporalis; PT, posterior part of temporalis; SM, superficial head of the masseter; DM, deep head of the masseter; MP, medial pterygoid; IP, inferior head of lateral pterygoid; SP, superior head of lateral pterygoid; AM, anterior mylohyoid; PM, posterior mylohyoid; GH: geniohyoid; AD: anterior part of the digastric.

## Notes

### Competing Interest Statement

The authors have declared no competing interest.

